# The effect of healthy aging on change detection and sensitivity to predictable sturcture in crowded acoustic scenes

**DOI:** 10.1101/2020.02.05.935817

**Authors:** Mathilde de Kerangal, Deborah Vickers, Maria Chait

## Abstract

The auditory system plays a critical role in supporting our ability to detect abrupt changes in our surroundings. Here we study how this capacity is affected in the course of healthy ageing. Artifical acoustic ‘scenes’, populated by multiple concurrent streams of pure tones (‘sources’) were used to capture the challenges of listening in complex acoustic environments. Two scene conditions were included: REG scenes consisted of sources characterized by a regular temporal structure. Matched RAND scenes contained sources which were temporally random. Changes, manifested as the abrupt disappearance of one of the sources, were introduced to a subset of the trials and participants (‘young’ group N=41, age 20-38 years; ‘older’ group N=41, age 60-82 years) were instructed to monitor the scenes for these events. Previous work demonstrated that young listeners exhibit better change detection performance in REG scenes, reflecting sensitivity to temporal structure. Here we sought to determine: (1) Whether ‘baseline’ change detection ability (i.e. in RAND scenes) is affected by age. (2) Whether aging affects listeners’ sensitivity to temporal regularity. (3) How change detection capacity relates to listeners’ hearing and cognitive profile. The results demonstrated that healthy aging is associated with reduced sensitivity to abrupt scene changes in RAND scenes but that performance does not correlate with age or standard audiological measures such as pure tone audiometry or speech in noise performance. Remarkably older listeners’ change detection performance improved substantially (up to the level exhibited by young listeners) in REG relative to RAND scenes. This suggests that the capacity to extract and track the regularity associated with scene sources, even in crowded acoustic environments, is relatively preserved in older listeners.

---

The ability to detect abrupt changes in our surroundings has serious, and sometimes immediate, implications for survival. The auditory system is hypothesized to play a key role in the brain’s change-detection network by serving as an ‘early-warning system’, continously monitoring the unfolding acoustic environment and rapidly directing attention to new events in the scene (Cervantes Constantino et al., 2012; Demany et al., 2010; Murphy et al., 2013). Indeed, sound is often what alerts us to important changes around us - in many cases we *hear* a change before we see it. Accumulating evidence demonstrates that (young) listeners are sensitive to abrupt changes, such as the appearance or disappearance of a source, even in heavily populated acoustic scenes (Cervantes Constantino et al., 2012; Pavani and Turatto, 2008; Petsas et al., 2016). Brain responses, recorded from naïve distracted listeners reveal that these events are often detected in the absence of directed attention (Sohoglu and Chait, 2016a, 2016b), consistent with an, at least partially, automatic change detection process. Here we ask how this critical capacity is affected in the course of healthy ageing.

Ageing is associated with loss of function within the peripheral and central auditory system that leads to a gradual loss of hearing ability (Presbycusis; Cruickshanks et al., 1998; Gates and Mills, 2005). Formally, emerging hearing difficulties are revealed by increased hearing threshold levels (usually measured with pure tone audiometry) and impaired speech comprehension, in particular in noisy environments (Dubno et al., 1984; Helfer and Wilber, 1990; Pichora-Fuller and Souza, 2003). A large body of work has focused on understanding the impact of healthy ageing on speech perception (Ben-David et al., 2012; Füllgrabe et al., 2014; Helfer and Freyman, 2008; Helfer and Wilber, 1990; Humes et al., 2012; Peelle and Wingfield, 2016; Pichora-Fuller and Souza, 2003; Tremblay et al., 2003; Zekveld et al., 2011). However relatively less is understood about how aging impacts the auditory system’s ‘early warning’ function, despite its key role in successful auditory scene analysis and profound implications to quality of life.

To capture the challenges of listening in complex acoustic environments, we used artificial acoustic ‘scenes’, populated by multiple concurrent streams of pure tones (‘sources’; Figure 1). These stimuli model crowded multi-object acoustic scenes akin to those we encounter during every day listening. Each ‘source’ had a unique frequency and rate commensurate with those characterizing many natural sounds. Changes, manifested as the abrupt disappearance of one of the sources, were introduced to a subset of the trials and participants were instructed to monitor the scenes for these events. Similar stimuli were previously used for the study of change detection ability in young listeners (Aman et al., 2018; Cervantes Constantino et al., 2012; Sohoglu and Chait, 2016a, 2016b).

**Figure 1:**
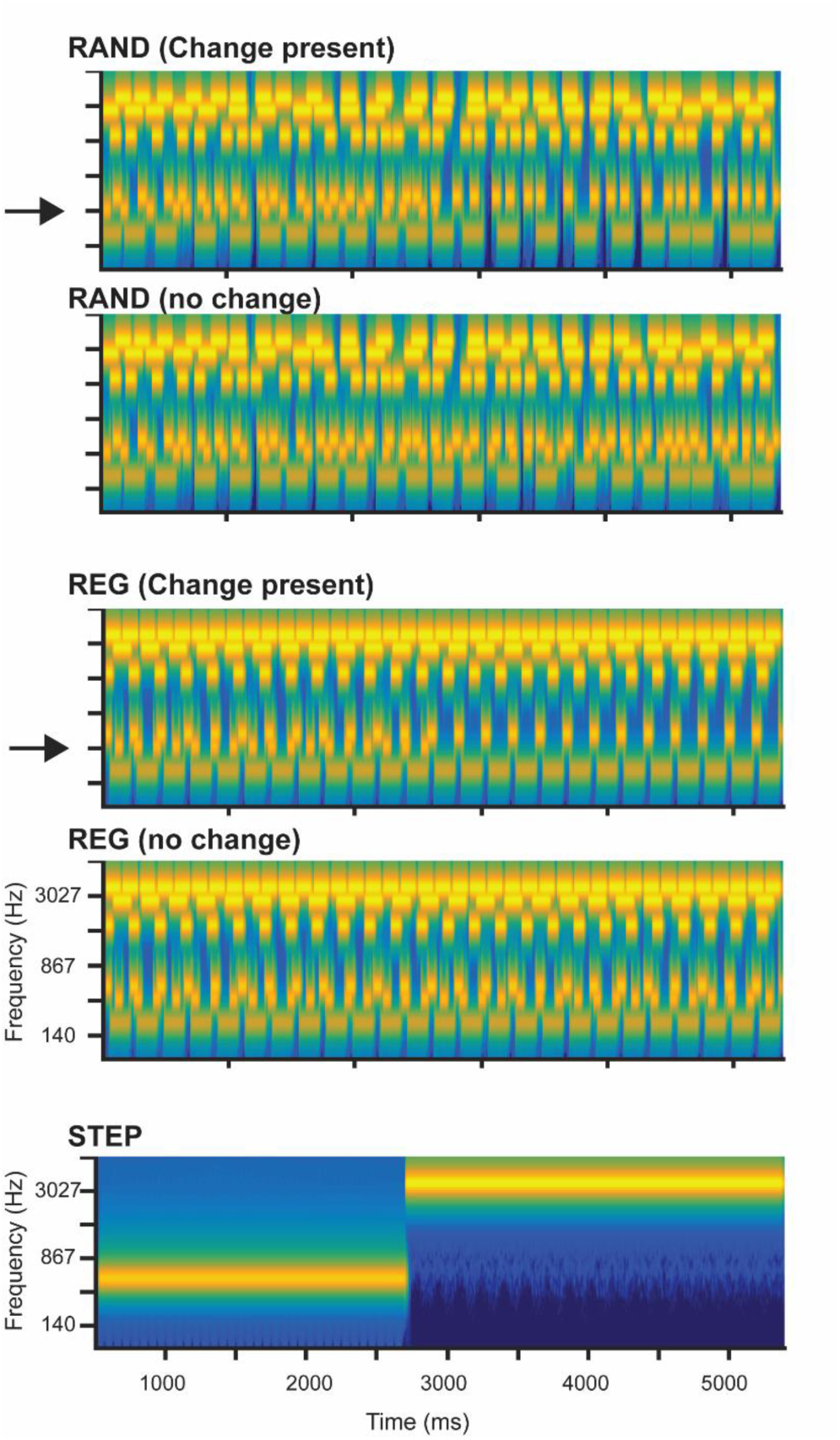
Stimulus examples. ‘Scenes’ consisted of six concurrent streams, each with a unique frequency and rate. Random (RAND) and regular (REG) scenes were matched spectrally, with only the temporal structure differing between conditions. All were presented in an intermixed, random order with 50% of the trials containing a change in the form of a disappearance of one of the scene sources. The disappearing components are indicated with arrows. Also plotted is an example of the control stimulus STEP. Participants were required to respond as quickly as possible to the scene changes and the frequency change in the STEP stimuli (the latter served as a control to estimate participants’ baseline reaction time). The overall stimulus duration ranged between 4 and 7 s, with the changes occurring between 2 and 3.5 s. The plots represent ‘auditory’ spectrograms, generated with a filter bank of 1/ERB wide channels equally spaced on a scale of ERB-rate. Channels are smoothed to obtain a temporal resolution similar to the Equivalent Rectangular Duration.

Whilst very simple, a major advantage of these stimuli is that they reduce reliance on semantic cues and allow for tight experimental control to isolate parameters of interest. Here, scene streams were set apart in frequency (at least 2 ERB; Moore and Glasberg, 1983) to reduce energetic masking and to facilitate relating individual audiological thresholds to change detection performance. Individual stream rates were between 3 – 25 Hz in line with the temporal properties which characterize many behaviourally relevant sounds including key rates in speech (Rosen, 1992). Previous work with young listeners (Cervantes Constantino et al., 2012) revealed high sensitivity to scene changes, despite the fact that changing components are not known in advance. The pattern of performance exhibited by listeners suggested that they rely, at least partially, on mechanisms which automatically track the individual scene sources and signal stream offset by generating a “local transient” (Cervantes Constantino et al., 2012).

Change detection ability also depends on sensitivity to regularity. Listeners perform substantially better in scenes which contain predictable, relative to unpredictable, temporal structure (Aman et al., submitted; Sohoglu and Chait, 2016b). This is in line with many previous demonstrations that observers are tuned to predictable patterning in sensory signals and use this information to efficiently interact with their surroundings (e.g. Andreou et al., 2011; Costa-Faidella et al., 2011; Lange, 2013; Leaver et al., 2009; Näätänen et al., 2011; Nelken, 2012; Rohenkohl et al., 2012; Winkler et al., 2009; Yaron et al., 2012). The regularity-linked improvement in change detection performance is hypothesized to reflect the fact that the auditory system forms models of the content of ongoing scenes. Novel events (e.g. appearance or disappearance of a source) that violate those models generate “prediction errors” which evoke greater neural responses and lead to increased perceptual salience (Sohoglu and Chait, 2016b). Sensitivity to source temporal structure is perhaps especially critical for disappearance detection - to efficiently determine source cessation, an ideal observer must ‘acquire’ the pattern of onsets and offsets associated with that channel, and respond as soon as an expected tone pip fails to arrive.

Despite mounting observations of the auditory system’s capacity to use predictable patterns to facilitate listening in crowded environments (e.g. Andreou et al., 2011; Bendixen, 2014; Bendixen et al., 2010; Winkler et al., 2009), the effect of aging on these processes remains largely unstudied. However, in a key report, Rimmele et al, (2012) provided some evidence of an age-related deficit in sensitivity to regular structure. Using a stream segregation paradigm, they showed that, unlike younger listeners, older listeners failed to take advantage of the regularity of an ignored stream to facilitate segregation. Here we seek to establish whether similar difficulties might generalize to a change detection task.

To determine whether older listeners are able to exploit predictable scene structure to facilitate change detection, we included two scene conditions (Figure 1): REG scenes consisted of ‘sources’ characterized by a regular temporal structure (specific rates were assigned randomly on each trial). RAND scenes were matched in terms of spectral structure, but contained sources which were temporally random. Based on previous observations in younger listeners (Aman et al., submitted; Sohoglu and Chait, 2016b) we expected that if older listeners are able to use temporal structure during scene analysis they should exhibit improved performance in REG relative to RAND scenes.

To focus on healthy aging, we sought to recruit participants with relatively preserved hearing and cognitive function. Forty-one participants (aged 60-85 years old) were recruited from the community of the university of the third age - a UK-wide movement which brings together retired people to pursue educational, social or creative activities. All had no, or at most mild hearing loss and demonstrated cognitive abilities within the normal range (see ‘inclusion criteria’, below). A control group of young listeners (N=41; between the ages 18-35) provided a performance benchmark against which the data from the older group were compared.

To ascertain whether change detection ability is predictable from the listener’s audiological status, we sought to correlate performance with standard measures of auditory function, including pure tone audiometry, measures of spectral resolution, and speech in noise processing. In particular, we aimed to understand whether change detection ability might draw on similar scene analysis mechanisms as those implicated in speech in noise processing (e.g. Holmes and Griffiths, 2019).

Beyond loss of hearing acuity, healthy ageing is also associated with various deficits of cognitive, executive and sustained-attention function which have expansive perceptual consequences across sensory modalities (Anstey and Low, 2004; Caserta et al., 2009; Harada et al., 2013; Juan and Adlard, 2019; Monge and Madden, 2016; Wetherell et al., 2002; Woodard and Sugarman, 2012). In the context of hearing, these deficits may have wide-ranging implications for listening in crowded environments. We therefore also collected performance measures on a battery of tests that capture broader cognitive abilities hypothesized to be affected by aging. Regression analyses were used to understand how listeners’ change detection ability might relate to more general age-related cognitive decline.

Overall, we address three main questions: **(1)** Whether ‘baseline’ change detection ability (i.e. in RAND scenes), is affected by age. **(2)** Whether aging affects listeners’ sensitivity to temporal regularity and **(3)** How change detection capacity relates to listeners’ hearing and cognitive profile.

## Methods

### Participants

#### Inclusion criteria

Three major inclusion criteria were used. Firstly, all participants were required to be in good general health, including no major surgery in the preceding two years. Secondly, participants were required to demonstrate cognitive abilities within the normal range. This was assessed using the ACE (Addenbrooke’s Cognitive Examination) mobile screening app (http://www.acemobile.org) delivered on an iPad (∼20 min). Lastly, we aimed to recruit participants with largely preserved hearing. This was formally defined as audiometric thresholds, in at least 1 ear, between -10dB – 30dB HL for frequencies between 125 – 2kHz and no worse than 40dB HL for 4 kHz. There were no requirements concerning the 6 - 8 kHz frequency range.

#### Groups

Two participant groups were tested. The **younger group** comprised 41 participants (32 females, age: 20 to 38, mean: 25.8) recruited from the UCL participant database. The **older group** comprised 41 participants (32 females, age: 60 to 82, mean: 69) who were recruited from the ‘University of the 3^rd^ Age” (U3A; https://www.u3a.org.uk/) and therefore represent a sample of high functioning older individuals.

In the older group, 20 participants had clinically normal hearing, defined as hearing thresholds <25 dB for the frequency range extending from 125Hz to 4kHz for both ears. The remaining older participants had at least one frequency between 125 Hz and 4 kHz with a threshold >25dB HL for at least one ear; these participants were considered to have mild hearing loss. Experimental procedures were approved by the research ethics committee of University College London and written informed consent was obtained from each participant.

### Procedure

The overall experimental session lasted approximately 2h30min (including breaks). Participants sat in a double-wall sound-proof booth and performed all assessments (including the main change detection task) in a random order, separated by breaks.

### Profile measures

In addition to the main experimental task (change detection) we administered a variety of tests to measure hearing performance and various cognitive abilities spanning the range of attention/memory/executive function, hypothesized to decline with healthy aging (Manly et al., 2000; Salthouse, 2011; Schoof and Rosen, 2014; Stuss et al., 1987; Tombaugh, 2004).

#### Audiometry

Audiometric thresholds were obtained for each participant for the following frequencies: 0.125, 0.25, 0.5, 1, 2, 4, 6 and 8 kHz. The audiogram was acquired using a GSI Pello audiometer presented over Radio Ear 3045 DD45 headphones. Two measures were derived for each participant: **LF Audiometric score** (mean threshold over the lower frequencies: 125hz to 4kHz) and **HF Audiometric score** (mean threshold over the high frequency range: 6-8 kHz).

#### Speech-in-noise test

A speech-in-noise comprehension score was obtained for each participant, using a coordinate response measure test introduced in Messaoud-Galusi et al, (2011). The participants were presented with a visual interface showing an image of a dog and a list of digits (1 2 3 4 5 6 8 9) and colours (black, white, pink, blue, green, red). They listened to a female speaker (target voice) producing the sentence: “show the dog where the [colour] [number] is”. The participants were instructed to tap on the correct combination and guess if unsure. The target voice was presented together with a 2 male-speaker babble that the participants were instructed to ignore. The babble consisted of understandable English sentences. The overall level was fixed at 70dB SPL. The SNR threshold between the babble and the target speaker was initially set to 20dB SPL and adjusted using a three-up one-down adaptive procedure to track the 79.4% correct threshold. The outcome SNR was calculated as the mean of the last 4 reversals. Each participant performed the test 4 consecutive times; the mean over the SNRs collected is used in the analyses below.

#### Spectral-temporally modulated Ripple test (SMRT)

The SMRT (Aronoff and Landsberger, 2013) uses spectral ripple stimuli to measure auditory spectral resolution. The test consists of an adaptive procedure whereby the ripple density of the target stimulus is modified until the listener cannot distinguish between the reference and the target stimulus. The discrimination threshold, expressed as number of ripples per octave, is used for the analyses below.

#### Auditory Sustained Attention (TEA)

The Test of Everyday Attention battery (TEA; Robertson et al., 1996) evaluates various facets of attention abilities in adults between 18-80 years of age. For the present study we selected one subtest of the TEA: the “elevator counting with distraction” task, designed to target sustained attention and ability to resist distraction. The test consists of streams of low- and high-pitch tones of which participants must count the low-pitch and ignore the high-pitch tones. Ten trials were administered overall. The resulting score was the number of correct trials.

#### Sustained Attention Response Task (SART)

The SART evaluates visual sustained attention ability. Participants monitor a serially presented visual stimulus sequence and are instructed to respond, by button press, to frequent non-target stimuli but withhold a response to infrequent target stimuli. The SART requires high levels of continuous attention and is sensitive to attentional lapses (Manly et al., 2000; Robertson et al., 1997). Here, a version of the SART as implemented in the Inquisit software package (https://www.millisecond.com/) was used. Stimuli were visual digits (0-9) with “3” designated as the “no-go” target. Outcome measures are %fail score (i.e. failure on no-go trials), and mean reaction time (RT) for correct “go” trials.

#### Trail Making test

The Trail Making Test (TMT) is a measure of mental flexibility, visual search ability, speed of processing, and executive function (Tombaugh, 2004). It is often used in neuropsychological test batteries because of its sensitivity to a variety of neurological impairments, including brain damage (Reitan, 1958) and dementia (Rasmusson et al., 1998). The test requires participants to connect (by sequential tapping) numbered circles arranged randomly on the screen. The outcome measure is the time taken to complete the sequence. In version A of the test, only number labels are present and participants must tap on them in ascending order. In version B, circles with letters and numbers are present and participants must alternate between numbers and letters (1-A-2-B-3-C …). Here we used a digital (iPad) version of the TMT (NeuRA; Neuroscience Research Australia). The (log) competition time (referred to as TMTa and TMTb, respectively) is used for the analyses presented below.

#### Musical training

The number of years of active musical practice (instrument or choir) was documented for each participant.

### Change detection task

The stimuli were artificial “soundscapes” designed to capture the challenge of listening in a busy acoustic environment in which many concurrent sound streams, each with a distinctive temporal pattern, are heard simultaneously (Eramudugolla et al., 2005; Snyder et al., 2012).

Stimuli (Figure 1) were 4000-7000 ms long artificial ‘scenes’ populated by multiple (6) streams of pure-tones designed to model sound sources. Each stream was characterized by a unique carrier frequency (drawn from a pool of twelve fixed values between 200 and 3453 Hz; spaced 2 cams on the ERB scale; Moore and Glasberg, 1983) and temporal pattern. In a previous series of experiments we demonstrated that these stimuli are perceived as a composite ‘sound-scape’ in which individual streams can be perceptually segregated and selectively attended to, and are therefore good models for natural acoustic scenes (Cervantes Constantino et al., 2012).

In the ‘regular’ scenes (REG), the duration of a tone pip (values uniformly distributed between 20 and 160 ms; 7 ms steps; each tone shaped by a 3 ms onset and offset ramp) and the silent interval between pips (values uniformly distributed between 20 and 160 ms; 7 ms steps) were chosen independently and then fixed for the duration of the scene so that the pattern was regular (see Figure 1, left column). This mimics the regularly modulated temporal properties of many natural sounds. In ‘random’ (RAND) scenes, tone duration was chosen randomly and remained fixed throughout the scene, but the silent intervals between successive pips varied (values uniformly distributed between 20-160 ms) resulting in an irregular pattern (See Figure 1, right column).

Scenes in which each source is active throughout the stimulus are referred to as ‘**no change**’ stimuli. Additionally, we synthesized scenes in which a source became inactive (disappeared) at some intermediate time during the scene. These are referred to as ‘**change present**’ stimuli. The timing of change varied randomly (uniformly distributed between 2000 ms and 3500 ms post scene onset). The identity of the changing component was chosen randomly from one of the following frequencies 405Hz, 708Hz, 1156Hz, 1819Hz, with a further constraint that it was not the lowest or highest carrier frequency in the scene. In REG scenes the nominal time of change was set to the offset of the last tone augmented by the inter-tone interval, i.e. at the expected onset of the next tone, which is the earliest time at which the disappearance could be detected. For disappearing sources in RAND scenes, it is impossible to define change time in this way (because there is no regular temporal structure). For the purpose of measuring reaction time (RT), the change time in RAND scenes was set to the offset of the last tone-pip augmented by the mean inter-pip-interval (90 ms). Because the distribution of inter-pip-intervals was identical in REG and RAND conditions, if temporal structure does not play a role in change detection, RT should be identical in both conditions.

The set of carrier frequencies and modulation patterns was chosen randomly for each scene, but to enable a controlled comparison between conditions, scenes were generated in quadruples so that each REG scene had its RAND equivalent, maintaining the same spectral structure. Similarly, each ‘change present’ scene had its corresponding ‘no change’ scene - sharing the same carrier frequencies and modulation patterns and only differing in the presence of a change. All stimuli were then presented in random order during the experiment, such that the occurrence of change (and its timing) were unpredictable. The inter-stimulus duration was set between 1 and 2.5s. A total of 240 scene sounds were presented to the participants: 120 RAND and 120 REG scenes in which 60 were ‘change present’ and 60 were ‘no change’ trials.

To obtain a measure of basic reaction time from each participant, the stimulus set additionally contained twenty STEP trials which were randomly interspersed between the other stimuli. STEP trials consisted of a continuous pure tone that changed in pitch (carrier frequency) partway through the trial. All STEP trials contained a change, which occurred randomly between 2 and 3.5 s. The overall duration of the STEP stimuli was set between 4 and 7s. The carrier frequencies of the step sounds were taken from the same pool of frequencies as the scene sounds.

The participants were asked to press a button as soon as they heard a disappearance in the scenes or a change in pitch in the STEP sounds. They first performed a short training block lasting 2-3 min and then completed the change detection task which was divided into blocks of 6-7min.

Stimuli were synthesized with a sampling rate of 44100 Hz and shaped with a 30 ms raised cosine onset and offset ramp. They were presented with a Roland Tri-capture 24 bit 96 kHz soundcard over headphones (Sennheiser HD 595) at a comfortable listening level (∼60-70 dB SPL), self-adjusted by each participant during the training block. Stimulus presentation was controlled using the *Cogent* software (http://www.vislab.ucl.ac.uk/cogent.php).

### Analysis

#### Profile measures

Depending on the shape of the distributions, we used t-tests for normally distributed data or non-parametric tests (Mann-Whitney test) for data that were not normally distributed. Correlation analyses were carried out to investigate the relationship between profile measures. Depending on the shape of the data, Pearson correlation, for normally distributed data, and Spearman correlations for non-normal data were used. The correlations were conducted separately for the young and older groups.

Due to the large number of profile measures, a factor analysis was carried out to understand how the measures grouped together and to generate factors that would be used in a regression analysis. The factor analysis was performed using Varimax rotation with Kaiser normalization. The data comprised the scores from the profile measures as detailed above (see also Fig 2). The factor analysis resulted in the extraction of 4 factors, following the Kaiser criterion (eigenvalues >1), the rotation converged in 5 iterations.

**Figure 2:**
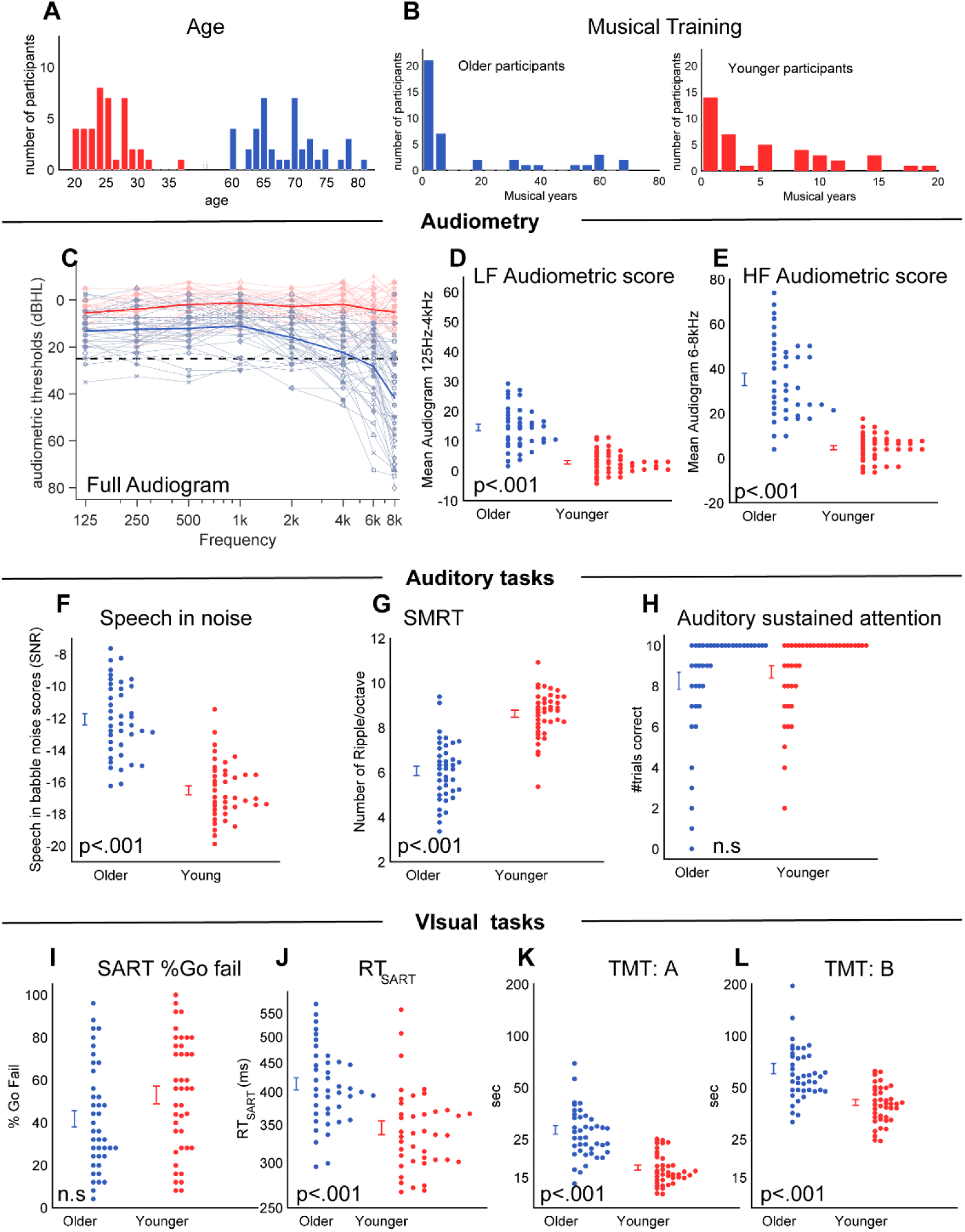
Summary of performance on profile measures; Red: young group Blue: Older group. As expected, the older group exhibited reduced performance, relative to the younger participants on most of the tasks, consistent with a growing hearing loss, and emerging aging related cognitive impairments **[A]** Age distribution **[B]** Distribution of musical training. The majority of participants were not musically trained as reflected by the greater number of participants on the left hand side of the graphs. **[C-E]** Pure tone audiometry scores (averages across ears). The dashed line in C marks 25dB which is commonly taken as the clinical threshold for the presence of hearing loss. The older listeners’ group exhibited overall higher thresholds in both the low frequency [D] and high frequency [E] ranges. **[F]** Speech in noise scores revealed impaired performance in the older group in line with previous demonstrations. **[G]** Performance on the spectro-temporally modulated rippled test revealed reduced sensitivity in the older group, in line with a growing impairment in processing rapid acoustic changes. **[H]** Performance on the Auditory sustained attention task (from ETA battery) resulted in ceiling performance across both groups. **[I-J]** Performance on the sustained attention to response task (SART) revealed a difference in reaction time between the two groups, but no difference in correct rate. **[K**,**L]** Performance on both versions of the trail making task yielded slower completion times from the older group indicative of impaired executive function.

#### Change detection task

Data from the change detection task were analysed in terms of Hit rate, False Positive rate, index of sensitivity d’, criterion C and reaction time (RT). d’, C and RT are used for the main statistical analyses. Sensitivity to regularity was quantified as the difference between the REG and RAND conditions. Parametric tests, t-test and repeated measures ANOVA, and non-parametric tests, Mann-Whitney and Kruskal-Wallis were used to analyse the data, as appropriate.

## Results

### Profile measures revealed expected differences between the young and older groups

Overall, the analysis of group differences (Figure 2) confirmed that, as expected based on the available literature, the older group exhibited differences in audiometric scores (Cruickshanks et al., 1998; Fitzgibbons and Gordon-Salant, 2010a), speech-in-noise scores (Schoof and Rosen, 2014), the trail making test (Salthouse, 2011; Stuss et al., 1987; Tombaugh, 2004) and in sustained attention (Manly et al., 2000).

#### Pure tone Audiometry

The audiogram was divided into low frequencies (from 125 Hz to 4 kHz), and high frequencies (6kHz and 8kHz). Means were calculated separately for each of these ranges and plotted in Figure 2 D, E. A repeated measures ANOVA with group and frequency range (LF, HF) as independent factors demonstrated an effect of group (F(1,80)=137.31, p<.001, partial η^2^ =.632), an effect of frequency range (F(1,80)=77.389, p<.001, partial η^2^ =.492) and an interaction (F(1,80)=54.923, p<.001, partial η^2^ =.407). Post hoc paired-sample t-tests revealed that a significant increase in audiometric thresholds with frequency was present for the older group (t(40)=-8.727, p<.001, Cohen’s d= - 1.363) but not for the younger group (t(40)=-1.867, p=.069, Cohen’s d= -0.292). These findings are consistent with a large body of work which chronicles a progressive loss of hearing acuity with age, which is particularly pronounced in the high frequency range.

#### Speech-in-noise test

The speech-in-noise scores are presented in figure 2F. An independent samples t-test demonstrated a significant group difference (t(80)=-9.8, p<.001, Cohen’s d= -2.171), consistent with previous observations (e.g. Schoof and Rosen, 2014).

#### Spectral-temporally modulated ripple test (SMRT)

The scores from the SMRT are shown in figure 2G. An independent samples t-test confirmed that the difference between older and younger groups was significant (t(80)=9.812, p<.001, Cohen’s d= 2.167). Older listeners exhibited higher ripples per octave thresholds, indicating reduced sensitivity to spectral information.

#### Auditory Sustained Attention

The results from the auditory sustained attention test are shown in figure 2H. Many participants, in both groups, were at or close to ceiling for this task. An independent samples Mann-Whitney test confirmed that the difference between the groups was not significant (U=791, p=.617, Cohen’s d= 0.187).

#### SART

Results from the SART test are shown in figure2 I, J. The %fail scores (Fig 2I) did not differ between groups (independent samples Mann-Whitney test, U=638, p=.061, Cohen’s d= 0.433). However, a significant group difference was observed for the RT measure (independent samples t-test, t(80)=5.09, p<.001, Cohen’s d= -1.124), showing, in line with previous demonstrations (Manly et al., 2000), slower reaction times for the older group.

We also tested whether the RT and %fail scores were correlated. Consistently with Manly et al. (2000), a significant correlation was found for both groups (Older: Spearman N=41, rho=-.733, p<.001; Younger: Spearman N=41, rho=-.743, p<.001) demonstrating that slower participants (longer reaction times) were also those generating fewer mistakes.

A second analysis was carried out to replicate a key finding from Manly et al. (2000) who showed that RT preceding successful no-go trials were significantly longer than those preceding failure to inhibit a response. We observed a similar effect: A repeated measures ANOVA with group and type of trial (preceding a correct or incorrect no-go trial) as independent factors demonstrated an effect of group (F(1,78)=30.8, p<.001, partial η^2^ =.282), an effect of type of trial (F(1,78)=57.3, p<.001, partial η^2^ =.423) but no interaction (F(1,78)=.019, p=.892, partial η^2^ <.001).

#### Trail Making Test (TMT)

The reaction time from TMT-A provides a measure of visual search speed and speed of processing for each participant. The TMT-B additionally provides a measure of “switching” or “mental flexibility. Both tests can also be considered as tests of general visual attention, as they require the participant to focus on an active visual task.

The TMT-A and TMT-B log reaction times are shown in figure 2K,L. A repeated measures ANOVA with factors group and condition (A or B) demonstrated an effect of group (F(1,80)=60.5, p<.001, partial η^2^ =.431), an effect of condition (F(1,80)=60.5, p<.001, partial η^2^ =.912) but no interaction (F(1,80)=1.07, p=.305, partial η^2^ =.013). In sum, the older subjects were generally slower than the younger group but the increased task difficulty in the TMT-B reduced performance to the same extent in both groups.

### No major cognitive differences between older listeners with mild hearing loss and those with clinically normal hearing

The older participants had good hearing overall but a subset fell below the clinical threshold for normal hearing (see methods). Based on these criteria a total of 21 older participants (50% of the cohort) were categorized as having mild hearing loss. With the exception of age (t(39)=-2.434, p=.020, Cohen’s d= -0.761), and (as expected from the literature) speech-in-noise performance (t(39)=-3.781, p=.001, Cohen’s d=-1.181), there were no group differences between the normal hearing older listeners (NH) and those with mild hearing loss in the remaining profile measures (Figure 3: musical training: Mann Whitney test U=185, p=.498, Cohen’s d= 0.046; auditory sustained attention scores: Mann Whitney test U=191, p=.603, Cohen’s d= 0.203; SMRT scores: t(39)=1.416, p=.165, Cohen’s d= 0.442; TMT-A: t(39)=-1.457 p=.153, Cohen’s d= - 0.455, TMT-B: t(39)=-1.665 p=.104, Cohen’s d= -0.520; SART %fail: Mann Whitney test U=187, p=.548, Cohen’s d= -0.144; SART RT: t(39)=1.920, p=.062, Cohen’s d= 0.600).

**Figure 3:**
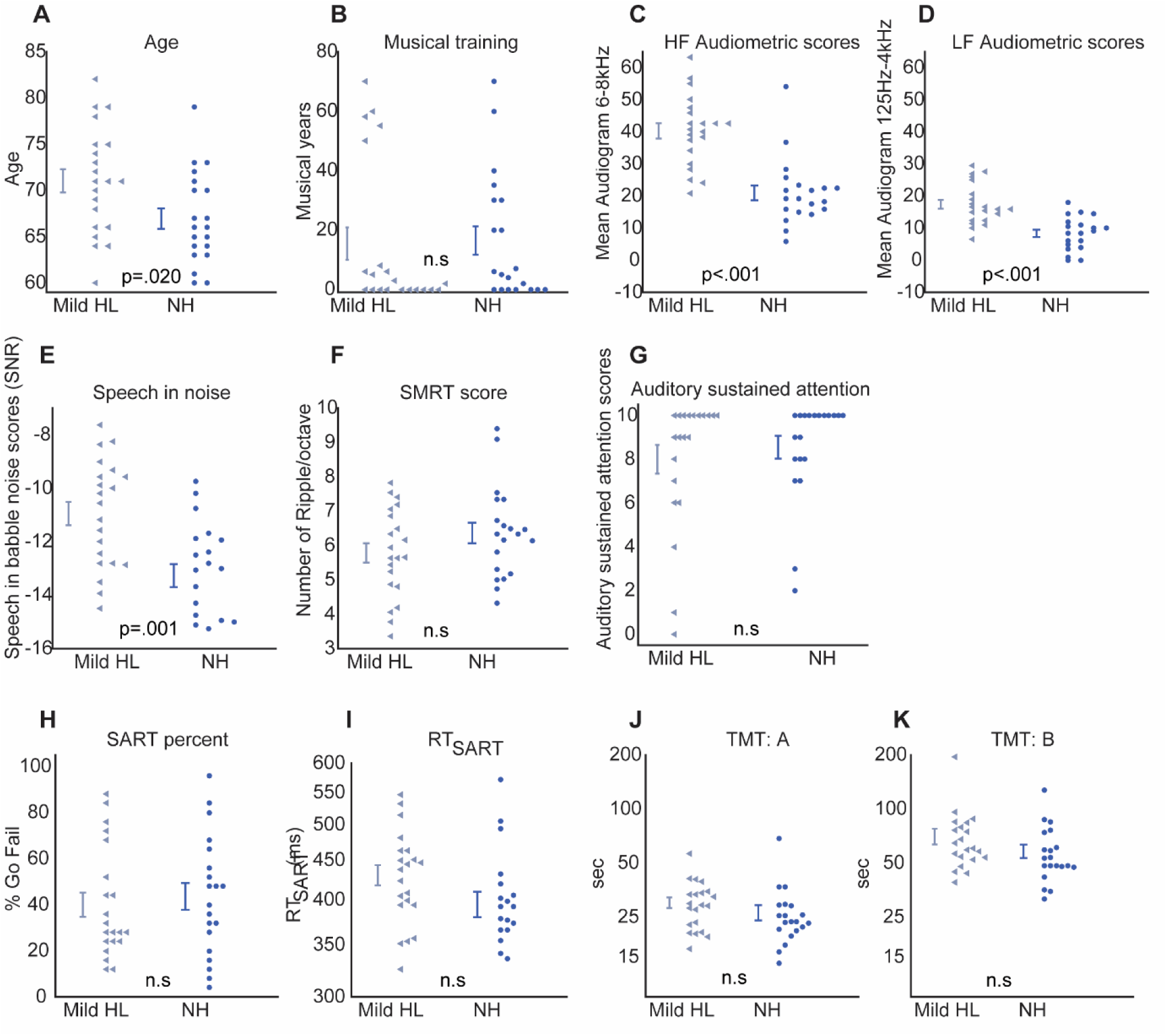
Performance on profile measures, separately for the normal hearing (NH; N=21) older listeners and those with mild hearing loss (Mild HL; N=21). The only differences observed were in age, pure-tone audiometry and speech in noise.

### Correlations between profile measures revealed a 4 factor latent structure

Correlations between profile measures were computed (for each age group separately) to identify potential links between the various cognitive abilities tested. Outcomes were not corrected for multiple comparisons and should therefore be regarded as exploratory in nature.

Hearing loss often progresses with age, and even though our older participants had relatively good hearing, a significant correlation was found between **age** and **LF audiometric score** (N=41, Pearson r=.404, p=.009) and **HF audiometric score** (N=41, Pearson r=.462, p=.002). Additionally, a significant correlation was found between **age** and **speech-in-noise** performance for the older group (N=41, Pearson r=.383, p=.014). These correlations were not observed for the younger group.

As reported in the literature (Moore, 2007), measures of hearing acuity can be related to speech-in-noise performance. Here, such correlations between pure tone audiometry and **speech-in-noise** performance were seen for the **LF audiometric score** (N=41, Pearson r=.506, p=.001) but not for the HF audiometric score. These correlations were not observed for the younger group.

Since sustaining attention on auditory information is an important component of speech-in-noise perception, we tested the correlation between these two measures; the measure of **auditory attention** correlated with **speech-in-noise** performance only for the younger group (N=41, Spearman rho=-.352, p=.024). Additionally, **TMT-B**, reflecting mental flexibility and visual search ability, correlated with **speech-in-noise** performance for both groups (Older group: N=41, Pearson r=.328, p=.036; Younger group: N=41, Pearson r=.340, p=.029). **TMT-B** also correlated with **SART RT** for the older group (N=41, Pearson r=467, p=.002). Older participants who were fast in the TMT-B tended to also demonstrate faster RT in the SART. No such correlations were observed for the young group.

**Years of musical training** correlated with the **auditory sustained attention** measure for the older group (N=41, Spearman rho=.512, p=.001) but not for the younger group.

Despite the SMRT scores differing between groups, they did not correlate with age or with any of the other profile measures in either group.

#### Factor analysis

Focusing on the older group only, a factor analysis was used to determine how the profile measures group together, understand the underlying structure of the correlation matrix and simplify the regression analyses (see below). The factor analysis resulted in the extraction of 4 factors, following the Kaiser criterion (eigenvalues >1), the rotation converged in 5 iterations.

Together, the extracted factors explained 68.09% of the variance in the data, with the first factor accounting for 21.02%, the second for 17.84%, the third for 15.3% and the fourth for 13.9%. The rotated factors are presented in table 1.

**Table 1:** factor loadings from the Factor analysis applied to the profile measures in the older group.

|  | Factor 1<br>Basic hearing<br>profile | Factor 2<br>SART | Factor 3<br>TMT | Factor 4<br>Auditory Sustained<br>attention |
| --- | --- | --- | --- | --- |
| Age | **.745** | -.100 | .156 | .230 |
| LF audiogram | **.765** | .113 | .008 | .181 |
| HF audiogram | **.761** | .162 | .147 | -.316 |
| Speech-in-noise | **.598** | .227 | .239 | -.213 |
| SMRT | **-.373** | .256 | .027 | .054 |
| SART scores | -.052 | **-.942** | .034 | -.082 |
| SART RT | .089 | **.866** | .231 | -.170 |
| TMT-A | .096 | -.071 | **.916** | -.022 |
| TMT-B | .177 | .284 | **.826** | -.072 |
| auditory sustained attention | -.146 | .161 | -.011 | **.803** |
| musical training | .161 | -.216 | -.079 | **.781** |

Consistent with the correlation results, the first factor mainly grouped measures of basic hearing ability-auditory acuity (LF audiogram, HF audiogram) speech-in-noise perception and SMRT. That age also loaded on this factor confirmed that these measures are tightly linked with aging. The second factor grouped the SART measures. The third was dominated by the measures from the TMT tests. The fourth factor grouped the measures of auditory sustained attention with musical training.

Overall the results of this analysis confirm that the profile measures used here captured 4 potentially separable cognitive dimensions. The first factor targeted basic auditory abilities (and age), the second and third were each dominated by a separate test, the TMT and SART, which apparently target at least partially distinct cognitive abilities in the visual modality. Perhaps the most unexpected was the fourth factor which explained a large share of the variance by loading musical training and auditory sustained attention.

These factors will be used for the regression analysis presented in the next sections.

### As a group, the older participants exhibited worsened ‘baseline’ change detection performance

The performance measured from the young cohort was used as a benchmark for evaluating change detection capacity in the older group. Firstly, we focused on understanding performance in the RAND condition. Subsequently we quantified any improvement arising from the introduction of regular structure (REG scenes). Data from one older participant were excluded from the analysis because of abnormally long reaction times.

The **hit** and **false positive (FP)** rates for both groups are shown in Figure 4A,B. A significant difference in hit rate was found between groups (Mann Whitney test, U=427, p<.001, Cohen’s d= - 0.848) with higher hit rates for the younger group. FP rates were generally low (mean in the older group=.0988, mean in the younger group=.0732), consistent with other change detection work (Cervantes Constantino et al, 2012; Aman et al., submitted), and did not differ significantly between groups.

**Figure 4:**
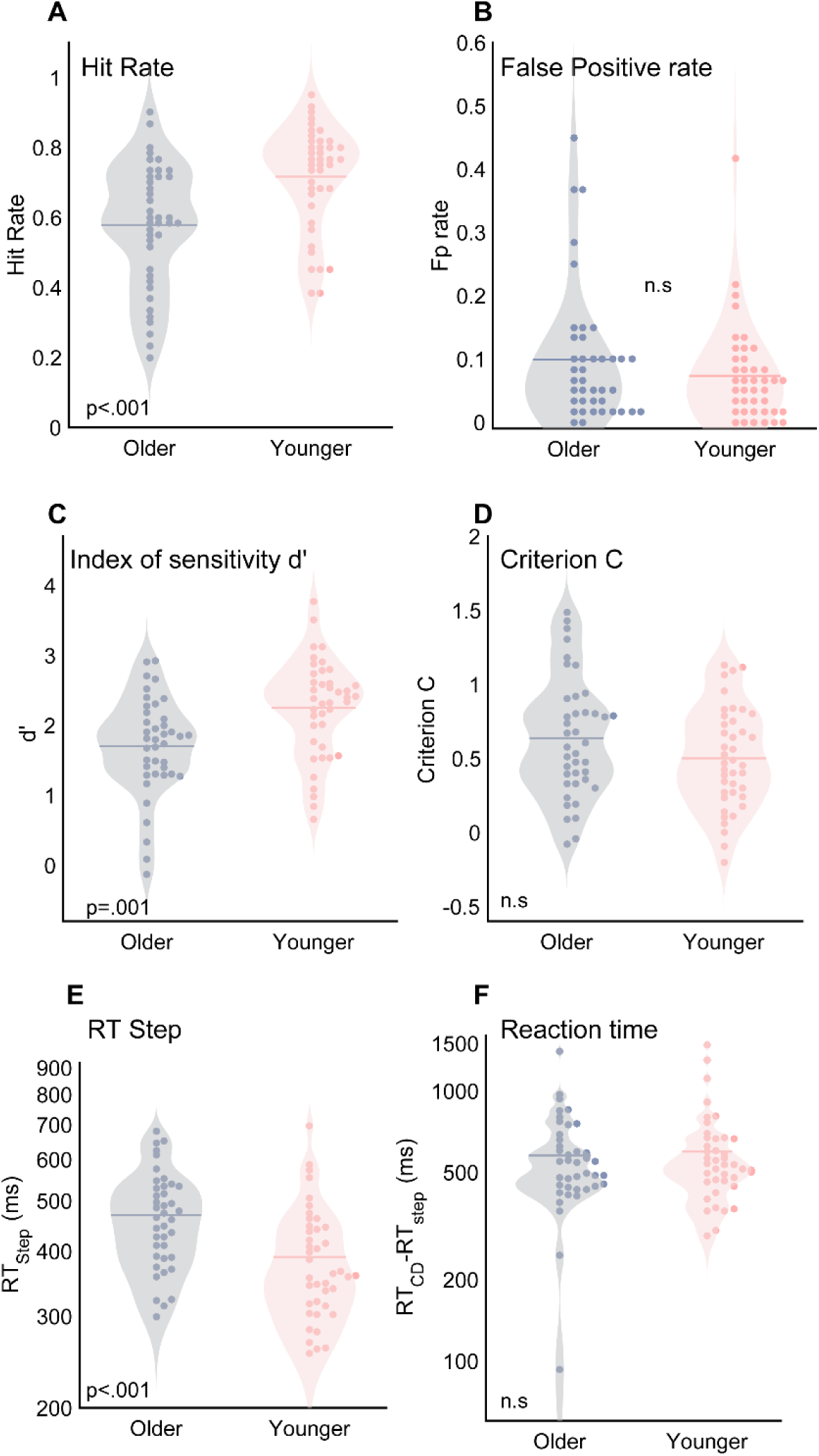
Performance on the RAND condition of the change detection task. This condition, where change detection can be achieved based on tracking changes in spectral scene content, is considered as the “basic” change detection condition. As a group, the older participants exhibited worse performance. Significant effects were observed on the hit rate and d’ measures. There was no difference in corrected RT between the older and younger groups.

The index of sensitivity **d’** and **criterion C** are measures derived from the hit and FP rates (Macmillan and Creelman, 2005). They are shown in figure 4C,D. The d’ values differed significantly between groups (Independent samples t-test, t(79)=-3.533, p=.001, Cohen’s d= -0.785), with higher d’ for the younger group. No difference was observed in criterion C values (Independent samples t-test, (t(79)=1.642, p=.105, Cohen’s d= 0.365).

**Reaction time (RT**; Figure 4E,F) was quantified by subtracting the RT to STEP sounds, which were interspersed between the scene change trials. The difference between groups for (log) RT was not significant (Mann-Whitney test, U=797, p=.828, Cohen’s d= -0.115). Younger participants were not faster than the older participants when responding to a disappearance, after the basic reaction time was accounted for.

The overall pattern of results demonstrates that older listeners’ reduced sensitivity to scene changes was driven primarily by reduced detectability (hit rate), with FP rates and bias not differing significantly from those of the younger listeners. In the following section we seek to understand whether the reduced performance (and increased variability) revealed by the older listener group is predictable from the profile measures.

### Change detection performance was largely independent of hearing ability

Firstly, we sought to determine whether the older listeners’ diminished performance was a consequence of impaired hearing acuity.

The carrier frequency of the disappearing component was selected from a set of 4 values: 405 Hz, 708 Hz, 1156 Hz or 1819 Hz. We correlated the hit rates associated with each of the disappearing component frequencies with the audiometric measure which was closest in frequency. The results are presented in table 2.

**Table 2:**
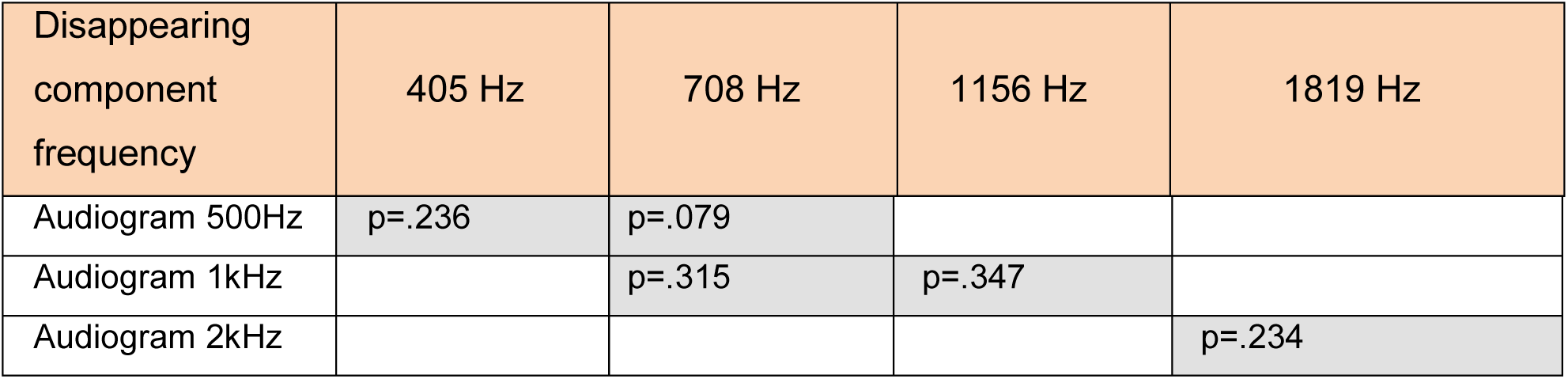
p values for the correlations between audiometric thresholds and change detection hit rates. The detectability of the change in each frequency range did not correlate with hearing acuity in that range as measured with audiometry.

There was no relationship between hearing acuity and Hit rate for the group of older participants tested. A further investigation of the effect of hearing acuity is below.

### Older listeners’ change detection ability is not correlated with standard audiological measures, but linked to sustained attention and history of musical training

A multiple regression analysis was conducted to understand which cognitive profile-derived factors correlate with change detection performance in the older group. The d’ based model significantly predicted 36.5% of the variance in the data (F(4,35)=5.028, p=.003). Only the 4th factor, dominated by musical training and auditory sustained attention scores, added significantly to the model (p<.001), while the first (p=.091), second (p=.764) and third (p=.145) did not. Similar analyses based on Criterion C and RT scores failed to significantly predict the data.

The correlation between the 4^th^ factor and d’ in RAND scenes is shown in Figure 5A: Those participants with high scores in auditory sustained attention/musical training also tended to demonstrate better change detection performance.

**Figure 5:**
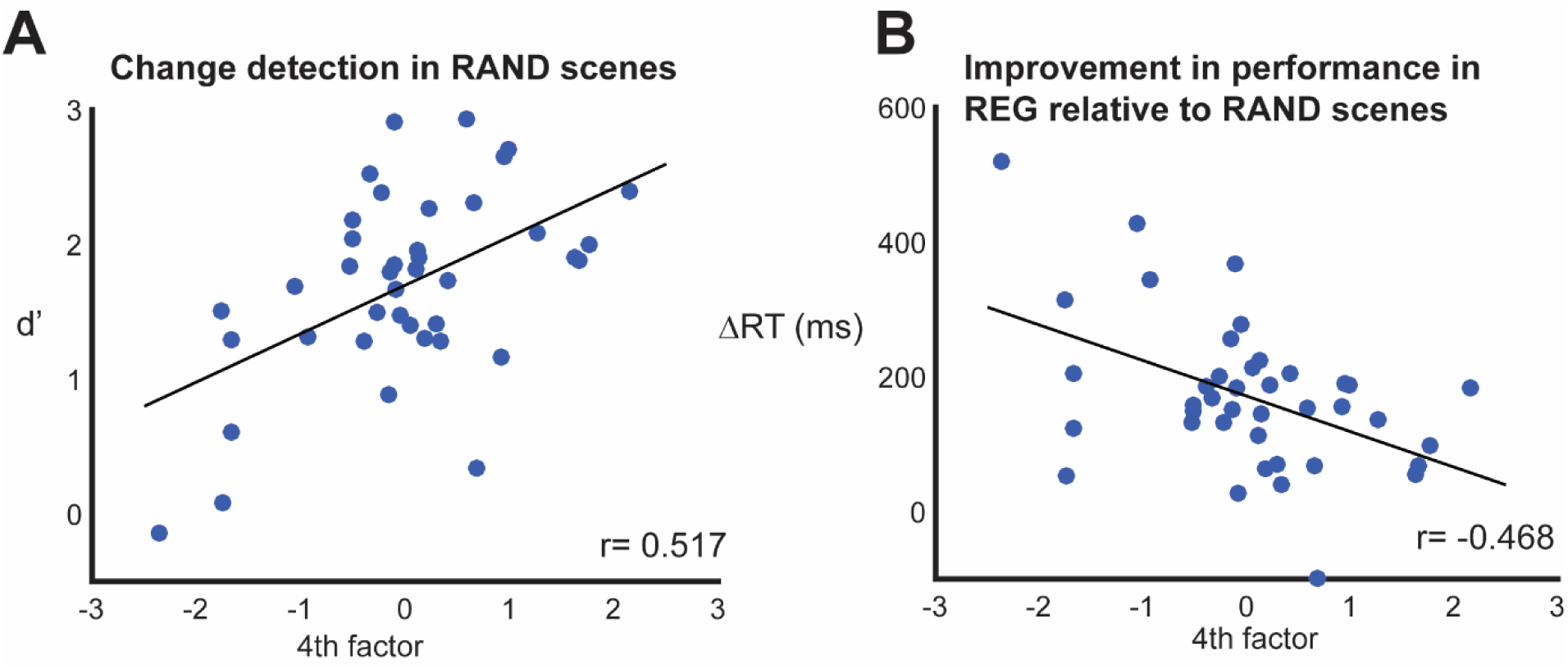
Older listeners’ change detection ability is linked to sustained attention and history of musical training. **[A]** Correlation (Pearson) between the 4^th^ factor, dominated by auditory sustained attention and musical training, and d’ in RAND scenes. This factor was identified in the regression analysis as the only set of profile measures which significantly explains d’ variability. High auditory sustained attention/musical training scores were associated with better change detection ability. **[B]** Correlation (Pearson) between the 4^th^ factor and ΔRT (difference in RT between REG and RAND scenes). The correlation indicates that those participants with the largest ΔRT were also those with lowest scores on musicality/auditory sustained attention.

### Older participants demonstrate retained sensitivity to regularity

Figure 6A presents the change detection performance (d’) in the random and regular conditions for all participants. A repeated measures ANOVA with group and scene condition as factors revealed a main effect of group (F(1,79)=7.686, p=.007, partial η^2^ =.089), main effect of scene condition (F(1,79)=200.54, p<.001, partial η^2^ =.717) and an interaction (F(1,79)=10.424, p=.002, partial η^2^=.117). Post-hoc tests confirmed that the interaction was due to the fact that performance of young and older listeners differed in the RAND scenes (t(79)=-3.533, p=.001, Cohen’s d=-0.785), but not in REG scenes (t(79)=-1.746, p=.085, Cohen’s d=-0.388). Older listeners were able to take advantage of the regular structure of scene components in the same way as did the younger listeners.

**Figure 6:**
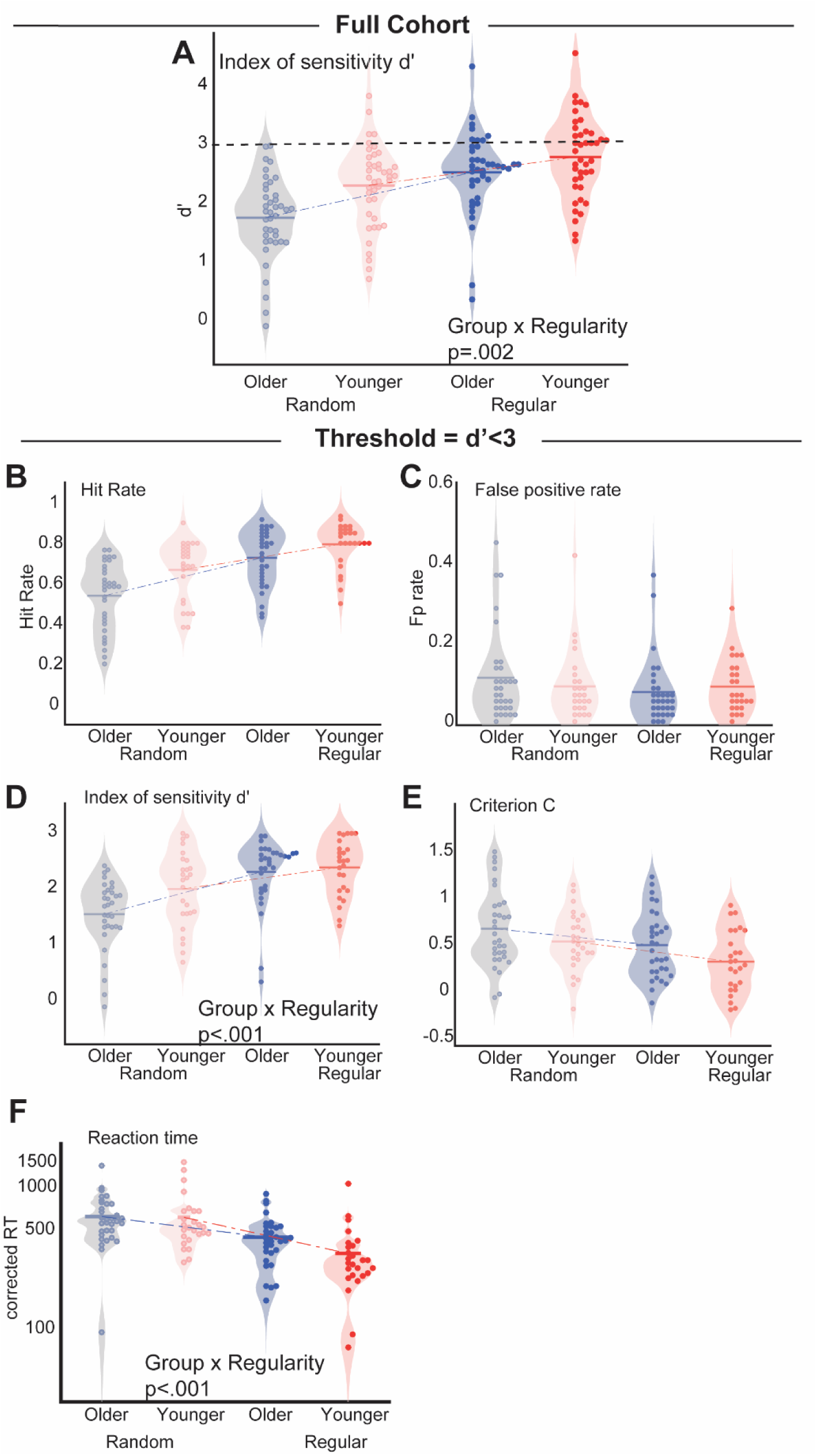
Comparing change detection performance in RAND and REG scenes. **[A]** Analysis of the full younger and older cohorts revealed a Group by Regularity interaction: whilst a difference exists between groups in RAND scenes (see also Figure 4), older listeners perform on par with the younger cohort in REG scenes. **[B-F]** change detection performance from a subset of participants whose performance fell below the ceiling threshold of d’=3 (dashed line in [A]). The interaction between Group and regularity persists, confirming it is not a consequence of ceiling performance

However, since many participants reached maximum performance in the REG condition, and particularly so in the younger group, there was a concern that ceiling effects might mask true differences in performance between groups. For the subsequent analysis (Figure 6B-F) we therefore focused on those participants who performed substantially below ceiling in all conditions (d’<3; see marked with a dashed line in Figure 6). The resulting cohort contained 26 young- and 32 older listeners.

Figure 6B,C presents the performance in random and regular conditions for hit rate and FP. With the introduction of regularity, the hit rates went up for both groups and the FP stayed stable. Hit and FP rates were used to calculate d’ and C on which the statistical analysis will focus.

Figure 6D,E shows d’ and criterion C in the RAND and REG conditions. A repeated measures ANOVA on d’ scores with group and scene condition as factors revealed an effect of condition (F(1,56)=133.2, p<.001, partial η^2^ =.704) and an interaction (F(1,56)=14.406, p<.001, partial η^2^ =.205) but no effect of group (F(1,56)=3.106, p=.083, partial η^2^ =.053). This confirms that the interaction observed for the full cohort held even with the high performing participants removed from the analysis. The d’ scores differed between groups in the RAND condition (t(56)=-2.665, p=.010, Cohen’s d=0.704) but not in the REG condition (t(56)=-.536, p=.594, Cohen’s d=-0.1415). The interaction reflects the fact that the difference between regular and random conditions was larger for the older participants than for the younger participants (difference between d’ REG and d’ RAND: t(56)= 3.796, p<.001, Cohen’s d=1.000). Namely, when the regularity cue was added the change detection performance of older participants improved more than that of the younger participants.

For the C results, a repeated measures ANOVA revealed an effect of scene condition (F(1,56)=28.981, p<.001, partial η^2^ =.341) but no effect of group (F(1,56)=3.293, p=.075, partial η^2^=.056) and no interaction (F(1,56)=.314, p=.578, partial η^2^ =.006). The introduction of regularity resulted in a similar decrease of C values for both age groups.

The reaction time data are shown in figure 5F. A repeated measures ANOVA demonstrated an effect of condition (F(1,56)=170.4, p<.001, partial η^2^ =.753) and an interaction (F(1,56)=16.256, p<.001, partial η^2^ =.225) but no effect of group (F(1,56)=2.280, p=.137, partial η^2^ =.039). The reaction times differed between young and older participants in the REG condition (t(56)=2.582, p=.012, Cohen’s d=0.682) but not in the RAND condition (t(56)=.189, p=.851, Cohen’s d=0.050). In the regular condition, the RT sped up in both groups, but to a larger extent for the younger listeners (difference between RT RAND and RT REG: t(56)=-4.032, p<.001, Cohen’s d=-1.065). An equivalent pattern is observed when analysing the full group of older participants (N=40). A repeated measures ANOVA demonstrated an effect of condition (F(1,79)=276.5, p<.001, partial η^2^ =.778) and an interaction (F(1,79)=17.09, p<.001, partial η^2^ =.178) but no effect of group (F(1,79)=.670, p=.415, partial η^2^ =.008). This suggests that though older participants appeared to benefit equally from regularity in terms of sensitivity (d’), they showed less improvement in RT relative to younger listeners.

Focusing on the older group, a multiple linear regression analysis using the profile measure factors as independent measures and d’ as the dependent measure significantly predicted 54.4% of the d’ variance (F(4,27)=8.067, p<.001). The fourth factor, dominated by auditory sustained attention and musical training, contributed significantly to the model (p<.001) while the others did not (1^st^: p=.272, 2^nd^: p=.667, 3^rd^: p=.342). Neither regression models for C or RT were significant (F(4,27)=1.215, p=.328; F(4,27)=.407,p=.802).

The benefit of regularity was quantified as Δd’= d’reg-d’rand, ΔC = Creg-Crand, ΔRT = RTrand-RTreg. A multiple linear regression using the factors as independent measures was performed to understand whether any of the collected profile measure can account for the benefit of regularity observed in the older listeners. For Δd’ (F(4,27)=1.080, p=.386) and ΔC (F(4,27)=1.050, p=.400), the predictions were not significant.

However the (log transformed) ΔRT based multiple linear regression significantly predicted 34.7% of the variance (F(4,27)=3.579, p=.018). The 4th factor contributed significantly to the model (p=.003). The contribution of the other factors was not significant (1^st^ factor: p=.955, 2^nd^ factor: p=.081, 3^rd^ factor: p=.850). As is seen in Figure 5B, the correlation indicates that those participants with the largest ΔRT were also those with lowest scores on musicality/auditory sustained attention. This outcome is consistent with the presence of a positive correlation between RT in the RAND condition and ΔRT (r=0.663; p<.001) and reflects the fact that those participants who found the RAND condition challenging, were also those demonstrating the largest RT improvement in REG scenes.

All of these conclusions held when running analyses on the full older cohort (N=40).

## Discussion

We demonstrated three main findings: firstly, as a group, older listeners exhibited worse performance than their young counterparts on the basic change detection task (RAND scenes). Second, sensitivity to regularity appeared largely preserved in the older listeners: the introduction of regular temporal structure (REG scenes) resulted in substantially increased performance - up to the same level measured in the young group. Third, older listeners’ change detection capacity did not correlate with age or any of the hearing-ability measures which we acquired (hearing acuity, speech perception in noise and spectral resolution). Instead it exhibited a strong correlation with auditory sustained attention ability and musical training. These results suggest that standard audiological assessments are not sufficient to predict age related deficits in auditory scene analysis.

### Basic change detection is impaired in older listeners

We used simple artificial acoustic scenes, devoid of semantic structure, to model listening challenges in crowded environments. RAND scenes (Figure 1), where each stream is characterized by a unique frequency and tone-pip duration, constitute the ‘basic’ condition. In these scenes, change detection is achievable by tracking spectral content. Older listeners exhibited a deficit in this condition, manifested as reduced hit rates (and d’) relative to the young group. Their false alarm rates, bias (criterion C) and (corrected) reaction times did not differ from the young group, suggesting a specific impairment in hearing out the disappearance event.

Correlation analyses suggested that older listeners’ reduced performance cannot be accounted for by hearing loss: Listeners’ general audiometric profile (1^st^ factor, See Table 1) which loaded age, pure-tone audiometry, spectral resolution and speech reception, did not correlate with change detection ability.

Instead, older listeners’ worse performance may be a consequence of reduced availability of processing resources leading to inability to simultaneously track multiple concurrent objects (Naveh-Benjamin et al., 2014; Peelle and Wingfield, 2016). Diminished resources may be a general consequence of aging (Glisky, 2007; Pronk Marieke et al., 2019) or result from an increased auditory perceptual burden that is not captured by standard hearing measures. It may also be the case that change detection ability is impacted by impaired temporal encoding of stream properties (e.g. Fitzgibbons and Gordon-Salant, 2015, 2010b, 2001), or impaired segregation of the concurrent streams (but see Alain et al., 1996; Snyder and Alain, 2006). Whilst future work is needed to pinpoint the source of the deficit, the outcome of the present study suggests that healthy aging is associated with a deficit in a critical auditory capacity and that standard measures of hearing function or speech processing may not be sufficient to diagnose it. We will return to this point later.

It should be noted that, for reasons which we detailed above, the ‘scenes’ used here are acoustically very simple. To understand how the impaired capacity identified in the lab setting extends to real life situations, it is important to link these measures with more complex naturalistic stimuli (e.g. Gregg et al., 2017; Pavani and Turatto, 2008).

### Older listeners demonstrate a largely preserved sensitivity to temporal regularity

Accumulating evidence, across sensory modalities, suggest that sensitivity to regularity plays a key role in shaping our perception of our surroundings (e.g. Andreou et al., 2011; Costa-Faidella et al., 2011; Lange, 2013; Leaver et al., 2009; Näätänen et al., 2011; Nelken, 2012; Rohenkohl et al., 2012; Winkler et al., 2009; Yaron et al., 2012). In contrast to previous reports (Rimmele et al., 2012), we found that older listeners exhibited a largely preserved sensitivity to temporal regularity. In REG scenes, their performance (d’) was identical to that of the younger group, including when controlling for potential ceiling effects. These results demonstrate that older listeners maintain the ability to track the predictable structure of ongoing sound input and register when it is violated, even when the scene is heavily populated with concurrent objects and the identity of the changing component is in advance unknown. Notably, however, the improvement in RT was smaller than that exhibited by the younger group, suggesting a potential residual difficulty in the older group.

The (largely) preserved sensitivity to regularity is consistent with other reports, e.g. from the beat synchronization literature which reveal that older listeners compensate for noisier temporal perception through reliance on predictive mechanisms (e.g.Turgeon et al., 2016; though the rates used in that research are usually substantially slower than those studied here). Similar effects are also commonly reported in the context of speech perception where older listeners are consistently demonstrated to compensate for hearing degradation by optimizing the use of the semantic and other predictive contexts (Lash et al., 2013; Nittrouer and Boothroyd, 1990; Peelle and Wingfield, 2016; Schneider et al., 2010; Schoof and Rosen, 2014).

Despite the conceptual similarity with that work, the present effects are likely tapping different, lower-level, mechanisms. The relevant predictive cues in our REG scenes involve rapidly unfolding information (over several concurrent streams) that precludes overt perceptual tracking and likely engages automatic statistical tracking mechanisms (Sohoglu and Chait, 2016b). Importantly, the rates used here– 3-25Hz – are all within the range that is considered to be most critical for hearing in natural environments (Kayser, 2019; Overath et al., 2015; Teng et al., 2017; Yi et al., 2019). That older listeners exhibited a benefit of regularity therefore indicates that the capacity to extract rapid temporal structure is largely maintained with healthy aging. Hearing aids should thus strive to preserve as much of this information as possible, to facilitate optimal auditory processing in crowded environments.

It is critical for future work to establish whether the conserved sensitivity to regularity observed here also translates to situations in which older listeners are passively exposed to the sounds without performing a task (e.g. as in Sohoglu and Chait, 2016a). Arguably, “change detection” is most useful as an “early warning” process, when we are not actively paying attention to the auditory scene.

Furthermore, the older people studied here represent a group of highly intellectually and physically active individuals. An important next step is to sample more broadly from within the aging population.

### Age-related effects on change detection are not correlated with standard audiological measures

In addition to the main change detection task, we collected performance on a set of commonly used general auditory- and cognitive-tests. The aim was to understand whether change detection ability can be predicted from an individual’s hearing and cognitive profile. As expected, the older group exhibited previously documented differences in audiometric scores, speech in noise perception, visual sustained attention ability, and executive function. A factor analysis (Table 1) suggested that the test battery captured 4 facets broadly related to **(1)** basic auditory function (including audiometry, spectral resolution, speech-in-noise processing ability and age) **(2)** General attentive vigilance, **(3)** Executive function, and **(4)** a factor that grouped sustained auditory attention task and musical training.

That the auditory- and visual-sustained attention tasks were linked with different components is consistent with the distinct demands associated with each test. The SART measured vigilance and propensity to inattention whereas the auditory sustained attention test tapped listeners’ ability to maintain attention on a target stream in the presence of a competing distractor.

That music and auditory attention loaded the same component suggests that those subjects who were musically trained also had better sustained attention ability and specifically in the auditory modality. However, as is commonly the case with such correlations, it is impossible to argue for a causal link.

Perhaps the most surprising outcome was that despite the overall impairment exhibited by the older group, a regression analysis revealed that standard measures of auditory function, including the ability to track a speaker amongst a competing babble, did not predict change detection performance. This suggest that notwithstanding potential similarities between speech-in-noise perception and change detection, these tasks capture different aspects of scene analysis. An important function of hearing in complex auditory scenes is therefore not adequately captured by current standard assessments resulting in sub-optimal understanding and management of age-related hearing degradation.

### A link between change detection and auditory sustained attention/musical training?

From amongst the present test battery, the factor that grouped auditory sustained attention and history of musical training was the only profile measure that consistently correlated with older listeners’ change detection capacity. It is noteworthy that the auditory sustained attention test used here (part of the TEA battery; Robertson et al., 1996) may not have been very sensitive to small differences between individuals – many participants, including in the older group, performed at ceiling (see Figure 2H). Our results suggest that developing more sensitive auditory sustained attention tasks (e.g. Zhao et al., 2019) may be crucial for accurate assessment of older listeners’ scene analysis ability.

Music training has been linked to enhanced performance on many listening tasks (e.g Bianchi et al., 2017; Parbery-Clark et al., 2011; Román-Caballero et al., 2018; Schellenberg and W. Weiss, 2013; Zendel and Alain, 2014). But whether the association is truly causal has been a matter of ongoing controversy. Regardless, the current results suggest that an individual’s life-long musical training background and sustained attention ability can predict their capacity to detect changes in complex auditory scenes.

More research is required to understand what the link between auditory sustained attention/musical training and change detection indicates about auditory processing in older listeners. That performance on the SART task did not correlate with change detection suggest that the underlying link is not related to vigilance per se, but rather aspects specific to listening capacity. For example, it may be that sustained attention/musical ability reflect the size of an individual’s “auditory processing bandwidth” or the availability of relevant computational capacity which is also required for change detection.

## Conclusions

The present work demonstrates that healthy aging is associated with loss of “listening ability” in complex acoustic environments that is reflected in reduced sensitivity to abrupt scene changes. The decline in such listening abilities is not well captured by standard audiological measures. Specifically, older listeners’ performance did not correlate with age, audiometry, or speech in noise performance. This suggests that change-detection-linked scene analysis abilities are independent from those associated with speech processing and audiologists and researchers should consider characterizing and assessing these aspects of auditory performance.

Remarkably, sensitivity to temporal structure appeared to be preserved in the older group: Older listeners’ change detection performance improved substantially (up to the level exhibited by young listeners) when temporal regularity was introduced. This suggests that the capacity to extract and track the regularity associated with scene sources, even in crowded acoustic environments, is relatively unaffected in aging.

## Acknowledgments

This project was supported by a PhD studentship from Action on Hearing Loss and a BBSRC project grant (BB/P003745/1) to MC. DV is supported by Medical Research Senior Fellowship (MR/S002537/1). We are grateful to Brian Moore and Stuart Rosen for advice and discussion.

